# Tandem gene clusters as phylogenetic anchors reveal the hidden history of vertebrate visual opsins

**DOI:** 10.1101/2024.02.06.579127

**Authors:** David Lagman, Christina A. Bergqvist, Shigehiro Kuraku

## Abstract

The expansion of the visual opsin gene family was a crucial event in the diversification of vertebrate vision in evolution. Additional expansions in phototransduction-related genes facilitated the development of dim-light (rods) and color vision (cones). Sequence-based phylogeny and gene positions from extant jawed vertebrate genomes are insufficient to untangle the visual opsin duplications in early vertebrates. Additionally, jawless vertebrates share a visual opsin gene repertoire with jawed vertebrates which conflicts with recent findings of distinct whole-genome duplications in each lineage. To resolve these questions, we analyzed jawless vertebrate genomes, focusing on visual opsin genes. Our findings, based on chromosomal arrangements and relationships, confirm tandem duplications of visual opsins before the vertebrate radiation.

## Introduction

Vertebrate visual diversity arises from differences in the absorption spectra of the visual opsins, light-detecting proteins used in the retina of the eye. These opsins, found in distinct photoreceptor cell types with related molecular architectures but some differences, contribute to visual capacities alongside the overall retinal organization. Consequently, the evolution of the visual opsins informs the origins of color vision and the emergence of different feature channels for visually guided behaviors in vertebrate evolution (Baden & Osorio 2019; Baden 2024).

Overall, five major types of visual opsins have been found in vertebrates: long wavelength-sensitive opsins (*LWS*), short wavelength-sensitive opsins (*SWS1* and *SWS2*), medium wavelength-sensitive opsins (*RH2*) and the dim-light rhodopsin (*RH1*) (Okano et al. 1992). The evolution of these genes have been suggested to have occurred through two scenarios (Fig. 1a and 1b) that must have taken place before the radiation of all vertebrates as evidenced by their identification in the pouched lamprey (*Geotria australis*) (Collin et al. 2003). The first hypothesis (Fig. 1a), based on sequence phylogeny, suggests a stepwise evolution of the different visual pigments (Okano et al. 1992), later suggested to be through tandem duplications followed by a whole genome duplication (Lamb & Hunt 2017). The second hypothesis (Fig. 1a), based on comparative synteny analyses in jawed vertebrate genomes, instead suggests a single tandem duplication generating two gene copies (an *LWS* ancestor and an *RH1/RH2/SWS1/SWS2* ancestor called OA in Fig. 1b) that then expanded in the early vertebrate whole genome duplications, called 1R and 2R, generating the five extant visual opsin subtypes (Lagman et al. 2013; Larhammar et al. 2009). Due to limited syntenic support in jawed vertebrate genomes for the first hypothesis and that it would require substantially more gene losses to account for the visual opsin gene numbers in extant vertebrate genomes we have considered it to be less likely than the second hypothesis. Especially, since sequence-based phylogenetic analyses are confounded by variable evolutionary rates, both among genes and across species (Kück et al. 2012; Kapli et al. 2021) and thus not always reflect the true sequence of duplications. Thus, in order to resolve the evolutionary history of visual opsins, analyses of conservation in many more species as well as representative vertebrate genomes among jawless vertebrates (diverging from jawed vertebrates ∼500 million years ago (Donoghue & Keating 2014)) is needed. To add another layer to the complexity of the situation, both hypotheses now has to be reevaluated in light of the recent evidence that jawed and jawless vertebrates only share 1R and that both lineages have experienced independent genome duplication (2R in jawed vertebrates) and triplication (CyWGD in jawless vertebrates) events respectively (Yu et al. 2024; Marlétaz et al. 2024; Nakatani et al. 2021). Although this complicates and invalidates the current hypotheses of the evolution of the visual opsin family, we could also use this as a tool to shed new light on the orthology and paralogy of visual opsins across all vertebrate lineages. Thus, the two central questions driving our current study are: 1) Do the genomes of jawless vertebrates support the first or the second hypothesis of vertebrate visual opsin evolution? And 2) Are the visual opsins in jawless and jawed vertebrates orthologous?

**Figure 1.**
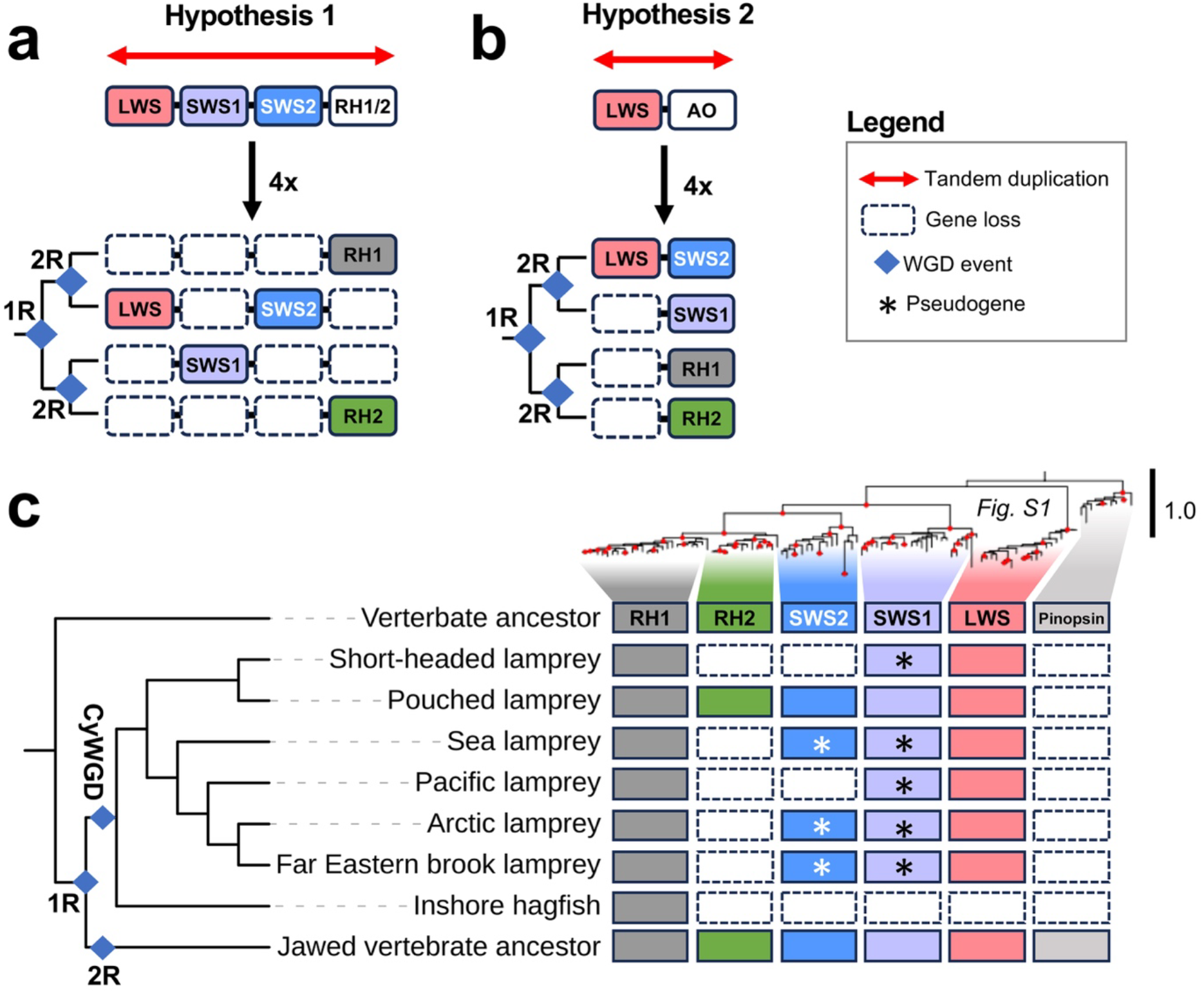
The phototransduction cascade and visual opsins. Different hypotheses proposed for visual opsin evolution in vertebrates (a and b). Scenario (a) is based on an analysis of the visual opsin molecular phylogeny (Okano 1992; Lamb 2022; Lamb & Hunt 2017) while scenario (b) relies on investigations into the visual opsin chromosomal regions of jawed vertebrates (Lagman et al. 2013). Cladograms next to chromosomes in (a) and (b) represent the proposed sequence of chromosomal duplications in these scenarios. AO (for Ancetral Opsin) in (b) represent an hypothetical ancestral opsin gene to *SWS1, SWS2, RH2* and *RH1*. c) Presence or absence of visual opsin genes in jawless fishes identified in this study combined with our updated molecular phylogeny (Fig. S1). The exon structure of the short-headed lamprey *RH1* gene, rearranged in the genome assembly, was curated based on transcriptome data to retrieve the full open reading frame. Filled red circles in the opsin tree indicate nodes with reliable supports (≥80% aLRT / ≥95% UFBoot support). The cladogram next to the opsin repertoires represent the evolutionary relationship between the jawless vertebrates and jawed vertebrates based on (Brownstein & Near 2023). 1R, the first WGD that was shared between jawed vertebrates and jawless fishes; 2R, the previously hypothesized shared second event; 2R, the jawed vertebrate specific WGD event; CyWGD, the jawless fish-specific hexaploidization event.

By analyzing all publicly available chromosome-scale genome assemblies of lampreys (sea lamprey, *Petromyzon marinus*; Arctic lamprey, *Lethenteron camtschaticum*; Far Eastern brook lamprey, *Lethenteron reissneri* and Pacific lamprey, *Entosphenus tridentatus*) we aimed to answer these questions. We also incorporated the newly sequenced genomes from two southern hemisphere species - the pouched lamprey, the only jawless vertebrate with a complete set of visual opsins (Collin et al. 2003) and the short-headed lamprey (*Mordacia mordax*) - to enable the first comprehensive comparison of opsin gene repertoires in lampreys. This analysis, with full recognition of their genomic locations, provides unprecedented evidence for reconstructing the history of diversifying opsin gene repertoires not possible previously.

## Results and Discussion

Sequence searches and phylogenetic analyses confirmed the visual opsin repertoire of the pouched lamprey, while several gene sequences could not be identified and some pseudogenization were observed in other species (Fig. 1c; see Fig. S1 for the full annotated tree based on exhaustive sequence sampling). The *RH1* gene in the short-headed lamprey had errors in the current genome assembly. This, in combination with a previous publication proposing *RH1* being a pseudogene (Lamb et al. 2016), prompted us to investigate transcriptome data. This effort led to the identification of the full-length coding sequence (Additional file 1), supporting its functionality.

Previous studies have observed that *LWS* and *SWS2* form a tandem pair in jawed vertebrates (same orientation, ∼10 Kbp apart in the spotted gar genome as an example), while the other genes are on separate chromosomes (Lagman et al. 2013). First we performed detailed searches in a selection of key jawed vertebrate taxa, in addition to the jawless species, (Supplementary table 1) and were unable to identify any other visual opsin combinations on the same strand in close proximity as would be required for confirmation of the first hypothesis. Validation of assembly completeness showed a low chance of false negative detection especially in jawed vertebrate species, considering high completeness scores for them (Figure S2). In order to confirm these findings in a larger set of vertebrates we extended the search for visual opsin gene clusters to include all vertebrate sequences in the NCBI RefSeq protein database. The identified visual opsin sequences were grouped into subtypes by phylogenetic inferrence (Fig. 2a; detailed clades in Figs. S4-S8; Supplementary table 2), then occurances of different subtypes found on the same chromosome and oriented similarly within 100 kbp or further apart, as a control, were counted. This analysis revealed that in jawed vertebrates, all instances of detected neighboring opsin genes within 100 kbp involve only *LWS* and *SWS2* genes (blue bar in Fig. 2b). Many of other detected instances, with longer intervals (orange bar in Fig. 2b), were from teleost genomes (∼88%, Fig. 2c) while the pairs with short intervals exhibited 50:50 representation of teleosts and non-teleost jawed vertebrates (Fig. 2c). This could most likely be explained by the frequent rearrangements observed in compact teleost genomes (Nakatani et al. 2007; Kasahara et al. 2007). In the lamprey genomes we observed two striking differences in the arrangement of visual opsin genes (Supplementary tabe 3). First our analyses reveal that the *RH2* and *SWS2* genes are adjacent to each other in the same orientation, approximately 6 Kbp apart in the pouched lamprey. Similarly, the *RH1* and *LWS* genes are positioned beside each other in the same orientation, less than 17 Kbp apart in all lamprey species except the Pacific lamprey where the *LWS* gene is divided between several scaffolds. Combined, these observations provide evidence for the first hypothesis (as suggested by Lamb & Hunt 2017), aligning with observations that tandem duplications play an important role in visual opsin gene multiplication of extant vertebrates (reviewed in Hagen et al. 2023).

**Figure 2.**
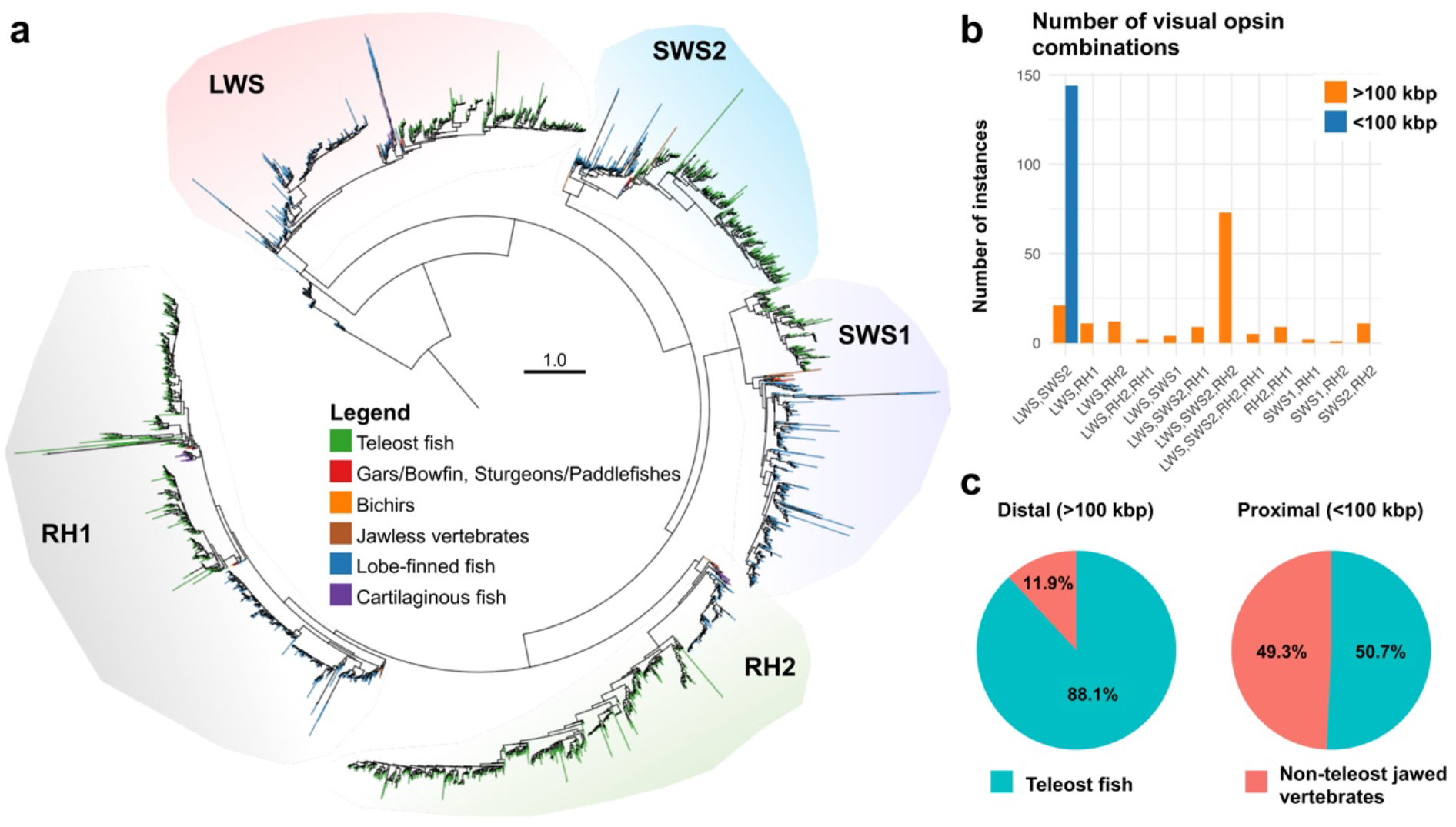
Vertebrate visual opsins identified in the RefSeq protein database and their subtype pairings on the same chromosome and strand. a) phylogenetic inference of ∼3300 vertebrate visual opsin sequences combined with the predicted jawless vertebrate sequences identified in this study. The full phylogeny was inferred using IQ-TREE with a constraint based on a reduced taxon set. Additional sequences were added and aligned with ClustalO prior to final inference. Node support values are omitted due to limited interpretability under the constrained analysis, particularly for newly added sequences. Branches are colored based on the groups of species listed in the legend. Detailed trees of each clade are presented in Figs. S4-S8 (LWS, SWS2, SWS1, RH2 and RH clades, respectively). The tree is rooted with OPN3 sequences. b) Counts of visual opsin subtype combinations present on the same chromosome and strand <100 kbp or >100 kbp apart in jawed vertebrate genomes. c) Percentages of teleost species and non-teleost jawed vertebrate species represented among these counts. Opsins counted were jawed vertebrate opsins from each subtype identified in the phylogenetic tree presented in (a).

Since jawless vertebrates most likely are ancient hexaploids (sharing 1R with jawed vertebrates followed by an independent genome triplication) we wanted to investigate if the visual opsin genes found in lampreys are orthologous to jawed vertebrate opsins. First, by phylogenetic inference on a concatenated alignment of genes flanking the visual opsins in lamprey genomes (Fig. 3a; Fig. S10; Supplementary table 4), we confirmed the orthologous chromosomes in all investigated lamprey species to sea lamprey chromosomes assigned to paralogon CLGE catalogued previously by Marlétaz et al. (2024). This analysis recapitulates the previous observations, with four of the clades forming two pairs (1B-1C and 2B-2C, respectively), separated in 1R. However, the placement of the two remaining clades (named 1A and 2A here) was not confidently supported in our analysis, likely due to the limited representation of some gene families on these chromosomes (Fig. 3a). The sea lamprey chromosomes represented in our 1A-1C clade was shown by Marlétaz et al. to be associated with spotted gar LG5 (which share a common ancestor with LG1 that duplicated in 2R according to their supplementary table 6). Thus by exclusion, the other sea lamprey chromosomes, represented in our 2A-2C clade, would be homologous to spotted gar LG3 and LG8. These chromosomal relationships of spotted gar chromosomes after 1R and 2R are also in agreement with (Lamb 2022). This indicates that the duplication generating the *RH1* and *RH2* genes indeed occurred in 1R (Fig. 3a and 3c) as suggested by Lamb & Hunt (2017). Additionally, we show that jawed and jawless vertebrates have retained *SWS2* genes that shared a common ancestor before 1R (Fig. 3a and 3c).

**Figure 3.**
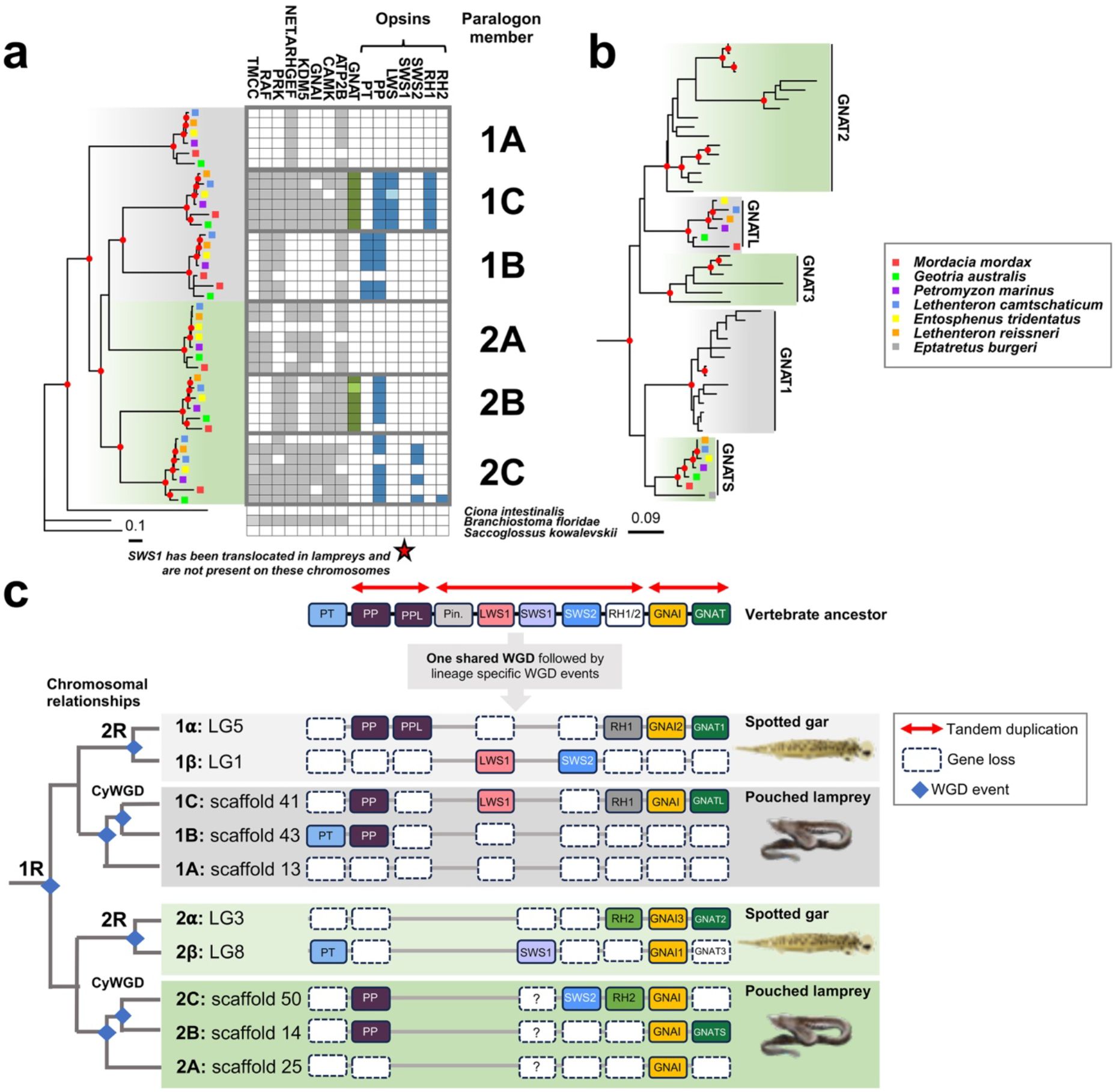
Evolutionary history of vertebrate visual opsins and GNAT genes. a) Molecular phylogeny of visual opsin-bearing chromosomal regions inferred from a concatenated alignment of genes flanking the visual opsins, including GNAI and parapinopsin (PP) (For detailed tree see Fig. S10). Grey and green shading is based on clade grouping in ref. (Marlétaz et al. 2024). Grey boxes depict the concatenated sequences per leaf, except PP. Colored boxes represent *GNAT* genes (green) or opsins (blue), including PP. Lighter shades represent sequences located on contigs or a different chromosome in the case of amphioxus (see Supplementary table 4). Star placed under the columns indicates the translocation of lamprey *SWS1* which have an uncertainty in which is the original chromosome. Clades are annotated with suggested lamprey paralogon member identifiers 1A-C and 2A-C. b) Molecular phylogeny of vertebrate *GNAT* genes (For detailed tree see Fig. S11). Tips without labels represent jawed vertebrate sequences. Grey and green shading represent chromosomal origin in 1R. The tree was rooted with the human GNAI amino acid sequences. In the molecular phylogenies nodes with reliable support (≥80% aLRT / ≥95% UFBoot support) are marked with filled red circles. c) Proposed evolutionary history of the visual opsins, their non-visual opsin relatives, and *GNAT* and *GNAI* genes in vertebrates. The cladogram illustrates the most parsimonious duplication scheme for these chromosomal regions in vertebrates. Chromosome/scaffold labels include paralogon identifiers, consistent with (a) for the pouched lamprey and ref. (Lamb 2022) for the spotted gar. VA opsin, another non-visual opsin, was excluded since it is in a separate genomic region. The *SWS1* gene was translocated in lampreys and the uncertainty in its original location has been indicated by a question mark. Additionally the pinopsin gene was translocated in Spotted gar and lost in lampreys. Abbreviated non-visual opsins are: PT – parietopsin; PPL – parapinopsin-Like; Pin. – pinopsin. Grey and green backgrounds represent origins of *RH1* or *RH2* carrying post-1R chromosomes/scaffolds, respectively. The spotted gar image was reused with the kind permission of Dr. Daniel Ocampo Daza (https://www.egosumdaniel.se).

Genomic regions harboring the visual opsins also contain the alpha subunit genes of the G-protein transducin (Larhammar et al. 2009; Lagman et al. 2013, 2012). In jawed vertebrates, rods and cones express *GNAT1* and *GNAT2* genes respectively while the sea lamprey have GNATS protein in rod-like photoreceptors and GNATL protein in cone-like photoreceptors (Muradov et al. 2008). Previously, lamprey *GNATS* and *GNATL* were considered orthologs of jawed vertebrate *GNAT1* and *GNAT2/3*, respectively (Lagman et al. 2012; Lamb & Hunt 2017). However, our updated phylogenetic analyses of the *GNAT* genes (Fig. 3b; Fig. S11), including lamprey sequences, revealed weak support for this orthology with the supports not reaching the confidence threshold (indicated by filled red circles representing ≥80% SH-aLRT and ≥95% Ufboot support in Fig. 3b). Additionally, our chromosomal synteny analyses suggest orthology to be the opposite to the previous conclusions (Fig. 3b and 3c). This finding aligns with the idea that gene family members have been reused multiple times, evolving similar biological systems/functions in various animal lineages (discussed in Oakley 2024) and also raises interesting questions and avenues of research about the origin of the rod and cone specific (with some exceptions) phototransduction cascades in vertebrates.

Our results reveal the complex evolutionary history of vertebrate visual opsins, characterized by tandem gene duplications, frequent losses, and differential paralog retention in jawed and jawless vertebrates. After the whole genome duplications (WGDs), few additional visual opsin genes persisted. Our new model (Fig. 3c) is based on comprehensive phylogenetic data, as well as exhaustive searches of tandem opsin genes as signals of phylogenetic reconstruction covering diverse jawless fishes.

## Materials and methods

Below follows a brief description of the methods, for detailed methods, see Supplementary methods provided in Additional file 3.

### Completeness of jawless vertebrate genomes

Completeness scores of jawless and jawed vertebrate genomes were calculated using BUSCO and compleasm with the odb10 ortholog set.

### Gene searches and phylogenetic analyses

Visual opsin and non-visual opsins as well as GNAT/GNAI sequences were searched in lamprey and selected jawed vertebrate genomes using TBLASTN. Automated gene prediction combined with manual annotation following sequence homology were performed. Sequences were classified via phylogenetic inference using alignments from Lagman et al. 2013 (opsins, teleosts excluded due to multiple tandem duplications) and from Lagman et al. 2012 (GNAT/GNAI).

To confirm the uniqueness of lamprey visual opsin subtype pairings (same strand and chromosome), BLASTP searches were performed against all vertebrate RefSeq proteins at NCBI. Opsins were identified through phylogenetic inference. Pairings were counted either proximal (<100 kbp) or distal (>100kbp) from other subtype genes. Sequences were aligned with ClustalO, and trees were inferred using IQ-TREE v3.

Sequence identities among visual opsins were calculated using all-versus-all BLASTP from the alignment used for Fig. 2a (Fig. S9).

### Gene family classification, selection, and analysis

Neighboring gene families to visual opsins in lampreys were identified. Protein sequences of vertebrate and outgroup species were grouped into gene families using OrthoFinder. Families with members on at least two sea lamprey chromosomes carrying visual opsins and present on at least three lamprey chromosomes were selected (seven families).

Peptide sequences from selected families were searched in lamprey genomes using TBLASTN. Automated gene prediction combined with manual annotation following sequence homology were performed. Lamprey sequences were combined with orthogroup sequences from vase tunicate, Florida lancet, and acorn worm. Sequences were aligned with ClustalO.

### Paralogon evolution

The sequences of selected neighboring family alignments were combined with alignments of GNAI and parapinopsins (PP) by chromosome/scaffold ID using the concatenate alignment function of MEGA 11. The resulting alignment was subjected to phylogenetic inference. Trees were inferred using IQ-TREE v3.

## Acknowledgements

We thank John Donald at Deakin University for early access to the unpublished genome assemblies, Mitsumasa Koyanagi and David Forsberg for commenting on the manuscript draft, Trevor Lamb and Dan Larhammar for insightful discussions, and Daniel Ocampo Daza for permission to use the illustration of the spotted gar.

## Data availability statement

The genome assemblies of the pouched- and short-headed lampreys are deposited in NCBI under BioProject PRJNA1030065. All sequence alignment files, and the phylogenetic inference output files have been deposited at FigShare (DOI: 10.6084/m9.figshare.29378324).

## Additional files

### Additional file 1

Nucleotide sequence of Short-headed lamprey rhodopsin from transcriptomic data

### Additional file 2

Supplementary tables 1-4.

### Additional file 3

Supplementary methods, supplementary figure legends and supplementary Figs. S1-S11.

## References

Baden T. 2024. Ancestral photoreceptor diversity as the basis of visual behaviour. Nat Ecol Evol. 8:374–386. doi: 10.1038/s41559-023-02291-7.

Baden T, Osorio D. 2019. The Retinal Basis of Vertebrate Color Vision. Annu. Rev. Vis. Sci. 5:177–200. doi: 10.1146/annurev-vision-091718.

Collin SP et al. 2003. Ancient colour vision: multiple opsin genes in the ancestral vertebrates. Current Biology. 13:R864–R865. doi: 10.1016/j.cub.2003.10.044.

Donoghue PCJ, Keating JN. 2014. Early vertebrate evolution. Palaeontology. 57:879–893. doi: 10.1111/pala.12125.

Hagen JFD, Roberts NS, Johnston RJ. 2023. The evolutionary history and spectral tuning of vertebrate visual opsins. Dev Biol. 493:40–66. doi: 10.1016/j.ydbio.2022.10.014.

Kapli P, Flouri T, Telford MJ. 2021. Systematic errors in phylogenetic trees. Current Biology. 31:R59–R64. doi: 10.1016/j.cub.2020.11.043.

Kasahara M et al. 2007. The medaka draft genome and insights into vertebrate genome evolution. Nature. 447:714–9. doi: 10.1038/nature05846.

Kück P, Mayer C, Wägele J-W, Misof B. 2012. Long Branch Effects Distort Maximum Likelihood Phylogenies in Simulations Despite Selection of the Correct Model Stiller, JW, editor. PLoS One. 7:e36593. doi: 10.1371/journal.pone.0036593.

Lagman D et al. 2013. The vertebrate ancestral repertoire of visual opsins, transducin alpha subunits and oxytocin/vasopressin receptors was established by duplication of their shared genomic region in the two rounds of early vertebrate genome duplications. BMC Evol Biol. 13:238. doi: 10.1186/1471-2148-13-238.

Lagman D, Sundström G, Ocampo Daza D, Abalo XM, Larhammar D. 2012. Expansion of Transducin Subunit Gene Families in Early Vertebrate Tetraploidizations. Genomics. 100:203– 211. doi: 10.1016/j.ygeno.2012.07.005.

Lamb TD et al. 2016. Evolution of vertebrate phototransduction: cascade activation. Mol Biol Evol. 33:2064–2087. doi: 10.1093/molbev/msw095.

Lamb TD. 2022. Photoreceptor physiology and evolution: cellular and molecular basis of rod and cone phototransduction. J Physiol. 600:4585–4601. doi: 10.1113/JP282058.

Lamb TD, Hunt DM. 2017. Evolution of the vertebrate phototransduction cascade activation steps. Dev Biol. 431:77–92. doi: 10.1016/j.ydbio.2017.03.018.

Larhammar D, Nordström K, Larsson TA. 2009. Evolution of vertebrate rod and cone phototransduction genes. Philos Trans R Soc Lond B Biol Sci. 364:2867–2880. doi: 10.1098/rstb.2009.0077.

Marlétaz F et al. 2024. The hagfish genome and the evolution of vertebrates. Nature. 627:811– 820. doi: 10.1038/s41586-024-07070-3.

Nakatani Y et al. 2021. Reconstruction of proto-vertebrate, proto-cyclostome and proto-gnathostome genomes provides new insights into early vertebrate evolution. Nat Commun. 12:4489. doi: 10.1038/s41467-021-24573-z.

Nakatani Y, Takeda H, Kohara Y, Morishita S. 2007. Reconstruction of the vertebrate ancestral genome reveals dynamic genome reorganization in early vertebrates. Genome Res. 17:1254–65. doi: 10.1101/gr.6316407.

Oakley TH. 2024. Building, Maintaining, and (re-)Deploying Genetic Toolkits during Convergent Evolution. Integr Comp Biol. 64:1505–1512. doi: 10.1093/icb/icae114.

Okano T, Kojima D, Fukada Y, Shichida Y, Yoshizawa T. 1992. Primary structures of chicken cone visual pigments: vertebrate rhodopsins have evolved out of cone visual pigments. Proc Natl Acad Sci U S A. 89:5932–6. doi: 10.1073/pnas.89.13.5932.

Yu D et al. 2024. Hagfish genome elucidates vertebrate whole-genome duplication events and their evolutionary consequences. Nat Ecol Evol. 8:519–535. doi: 10.1038/s41559-023-02299-z.

