## Supplementary material for "Tandem gene clusters as phylogenetic anchors reveal the hidden history of vertebrate visual opsins": Additional file 1.pdf

### **Additional file 1:** Short-headed lamprey rhodopsin (*RH1*) coding sequence

```
>c117566_g1_i1 Mordacia mordax RNA-seq assembly v1 len=2047 path=[1:0-83 85:84-84
86:85-504 5479:505-506 5482:507-1201 @1203@!:1202-1635 4843:1636-1639 1637:1640-
1746 1744:1747-2046]
CCGCCAATTAGACCCTGTGCTGCCACACAGCCACAAAGCTACCCCTAAGCCGACCCTCCAACCCTCCCTCCACGGGCGGG
GAGAGGGGGGGGGATTCCCCAGATGTCGCCCGGATCTTCTGGTTGCCGGTCCCAGGTTCCTCAAGCCGGGGAGACCTGGC
TGGTGGACACCGAGGACACCTCCGTCTTGCTCGTGGAGGCTCCCGAGTCATCGTCGCCCAGAGGATTCTTGCCGCAGCAC
AGCGTGGTGATCATGCAGTTGCGGAACTGCTTGTTCATGAGGATGTAGATGACCGGGTTGTAGAGGGACGAGCTCTTGGC
AAAGAAGGCCGGCGCGGTCATGAAGGTGGCGCCGAAGTCGGACCCCTGGTTGGTGAAGATGTAGAACGCCACGGAGGCGT
AGGGCACCCAACACACCAGGAATCCCACCACCATCAGTACCACCATGCGGGTCACCTCTTTCTCGGCCTTCTGCGTCGAC
GCCGACTCTTGCTGGGCCGCGGCCGCCTCCTTGACTGTGCAGAGCAAGCGTCCGTAGGAGAAGAAAATGATCACGAAAGG
GATGACGAAGTGTACGACGAACATGTAGATGACGAACGACTCGTTGTTGAAGTCGGGGTTCATCGTGTAGTAATCCGGCC
CGCACGAGCATTGCATTCCTTCCGGAATGAACCTGGACCAGCCTAGGAGTGGCGGAGCAGCGCATGACAAGGCCATGATC
CACGTGAAAGCCACCCCCATGATGGCGTGGGTGCTGCCGAAGCGGAAGTTGCCCATGGGTTTGCAAATGACGATGTAGCG
CTCAATGGCGAGCGCCACAAGGGACCAGAGAGACACTTCGCCGCCGAGCGTGGCAAAGAAGCCCTCCGTGGAGCACATGG
TGGGTCCGAAGATGAAGTAGCCGTTCATGGAGGTGTACATGGTGACGGTGAAGCCGCAGCAGATCATGAAGAGGTTGGAC
ACGGCGAGGTTGAGCAGGATGTAGTTGAGCGGGGTCCTCAGCTTCTTGTGCTGCACCGTGACGAACAGCGTGAGGAAGTT
GATGGGGAAGCCGACGAGGATGAGGAAGAACATGTAGGCAGCCAGGGCGGAGAACTTCCATGGTTCGGCCAGGTAGTACT
GCGGGTACTCGAAAGGACTGCGAACCACTCCGGTCTTGTTTGAGAACGGGACGTAGAAATTCTGTCCCTCTGTGCCGTTC
ATGGTGCTTGCTGTTGCCGAGTTCCCTGGTGCGTGCGGTGGTGAGATTTAACCAATGGGCACAAGTAGACGTGTGTGTTT
GCGATGTCTCCCGTGTGGATTGAATTCGTAAGTATTGCCTTGTTGTCGAGTGATTTGCTTAAAATGTTTTTTTTTTTAAAG
TCGTTTAACGGTTCAATTGAACAATTTGTTTTGATATTGGTAGTGGGTAGTGATGATACGTGTTGTGGTTTAAAAGCGTG
ATTAGTTCCAGTCGCGCCCACTTTGCCAAGAGTCTGATCGTGCGAGAGATGGCTTTCGCGCTGATGAAATAACTATAAAG
TGTTCATCAGAATAGGCGCTCGCTAACAGCAGTCGTCCACAGCAGACTTTATGAAGGCTGATGGCTCGATTGTGTTAACG
TGAATTAATGTGTACCACGAAGACGCAAGTGGACTGTGTTTGTCATGTGTTCATGTACACATGTGATGTAGGCGCACTAA
ACTTGTGTGAACTTGACGTTAATGTAATATCTACGTGTGTCGCTAATGTCCAGTCATGTGTGCACTGCGGTGGTGTTTCA
GTTAAGTGATGATCAACGGTATCTAGAGTTCTATGCTGTGCAGACAAGCGCTCGCTCAAGCGTTCGATACGGTGCGCGAT
ATGCAACGGCCGGTCGCTGGAATATACATAAATTCTGCACGACTGTGTAATCGTGTAGCTTGGGCACGTGGCGCGTTTGC
CAGAGCGTTAAATTGGATTCGCGCGTTCGTGGGTGCAAGGAAATGCCCTTGAGTTAAGCGATGACTGATGGCAGTGGCGC
ACGCGCCGTGCGGGTGCCGTGGTGCTGGTCCGTTCAGCAAGGGGGGG
```
