## Supplementary material for "Tandem gene clusters as phylogenetic anchors reveal the hidden history of vertebrate visual opsins": Additional_file_3.pdf

### Supplementary File 1

#### Supplementary methods

***Completeness assessment of genome assemblies***

Completeness of the genome assemblies used for exhaustive gene search were assessed primarily with BUSCO v5.8.2 (Manni et al. 2021), which were supplemented with assessment by compleasm v0.2.6 (Huang & Li 2023) when computation with BUSCO did not properly finish (indicated with asterisks in Figure S2). These assessments were performed using the odb10 ortholog sets prepared for individual taxa.

***Gene searches and phylogenetic analyses***

Visual opsin sequences from the pouched lamprey (*Geotria australis*), non-visual opsins from zebrafish (*Danio rerio*) and chicken (*Gallus gallus*), and *GNAT1*and *GNAI1* sequences from human were used as TBLASTN (Altschul et al. 1990) queries in detailed searches of genome assemblies from the pouched lamprey, short-headed lamprey (*Mordacia mordax*), sea lamprey (*Petromyzon marinus*), Arctic lamprey (*Lethenteron camtschaticum*), Far Eastern brook lamprey (*Lethenteron reissneri*), Pacific lamprey (*Entosphenus tridentatus*), inshore hagfish (*Eptatretus burgeri*), and several jawed vertebrate genomes that have been poorly studied with regard to visual opsin gene arrangemenets and representing key phylogenetic positions in jawed vertebrate evolution (see Supplementary table 1). The high sequence conservation among visual opsin subtypes (Fig. S9) and between *GNAT* and *GNAI* subtypes (Lagman et al. 2012) enabled reliable identification of genes within each lineage.

For each query, the top five hits were retained. High-scoring pairs (HSPs) within 50 kb were merged using bedtools merge (Quinlan & Hall 2010). Corresponding FASTA sequences were extracted using bedtools getfasta, and amino acid sequences were predicted with GeneWise (Birney et al. 2004) using default settings and the original BLAST query as a reference. The longest predicted peptide sequence per gene was retained for phylogenetic analysis.

Identified opsins were added to the visual opsin alignment from Lagman et al. (2013), excluding teleost species (due to multiple tandem duplication events in this lineage). *GNAT/GNAI* sequences were added to alignments from Lagman et al. (2012). For sequence information regarding the sequences used from these papers we refer to the respecitve supplementary tables with sequence information. Sequences predicted in this paper have sequence information in their sequence names otherwise we refer to the respective alignments for each sequence. Erroneous sequences were corrected by introducing gaps or through manual curation based on sequence homology to othr vertebrate sequences or RNA-seq data. All sequences were aligned using ClustalO (Sievers et al. 2011) with default parameters. Phylogenetic trees were inferred using IQ-TREE (v. 3) (Thi Hoang et al. 2017; Wong et al. 2025; Kalyaanamoorthy et al. 2017) with the following settings: -t BIONJ -nt 8 -quiet -keep-ident -bb 10000 -alrt 10000 -m TEST -bnni. Trees were visualized using iTOL (Letunic & Bork 2021), or in R using ggtree (Xu et al. 2022), treeio (Wang et al. 2020), and ape (Paradis et al. 2004).

Lungfish and inshore hagfish genomes were only searched for visual opsins to identify any paired subtypes beyond the previously known LWS–SWS2 pair in jawed vertebrates. Hagfish genomes were excluded from conserved synteny analyses due to extensive gene loss and reduced gene repertoires (Marlétaz et al. 2024).

To confirm the novelty of visual opsin gene pairings observed in lamprey genomes (LWS–RH1 and SWS2–RH2), we performed remote BLASTP searches using pouched lamprey visual opsins against the NCBI RefSeq protein database with the following parameters: -db refseq_protein -remote -entrez_query "Vertebrata[Organism]" -max_target_seqs 10000 -outfmt 6. Resulting hits were used in a secondary BLASTP search against chicken RefSeq proteins (NP_001384426.1, NP_990821.1, NP_990769.1,

NP_990848.1, NP_990771.2) with the following settings: -max_target_seqs 5 -num_threads 8 -outfmt 6. Sequences with at least one of the chicken visual opsins among the top hits were retained. Gene IDs, chromosomal positions, orientations, and amino acid sequences were retrieved from NCBI. The longest peptide sequence per gene was aligned using ClustalO, and a preliminary phylogenetic tree was inferred with IQ-TREE (v3.0.0) using the settings: -t BIONJ -nt 8 -quiet -keep-ident -m LG+G4 -fast. This allowed us to identify and remove non-visual opsins.

Due to the relatively short length of visual opsins (~300–400 amino acids) and the large number of sequences (>3000), reliable phylogenetic inference was limited by the low number of informative sites. To address this, we used CD-HIT (v. 4.8.1) (Li & Godzik 2006) to reduce redundancy at a 90% identity threshold. Predicted lamprey visual opsins were then added to the reduced alignment, resulting in a dataset of 787 sequences. A phylogenetic tree was inferred using IQ-TREE v3.0.0 with the following settings: -quiet -m TEST -T AUTO -B 10000 -alrt 10000 -nm 30000 -bnni.

This tree served as a backbone for re-integrating the sequences excluded by CD-HIT. These were added to the reduced alignment using ClustalO with the command: -i removed_sequences.fasta --p1 reduced_alignment.fasta. Final phylogenetic inference (Fig. 2a and Figs. S4–S8) was performed using IQ-TREE (v3.0.1) with the following settings: -g reduced_alignment.fasta.contree -pre combined_tree -m JTT+F+G4 -fast -alrt 1000 -T 8. Subtype assignments from the full tree were combined with gene positional data to identify co-located genes on the same chromosome and strand within or beyond 100 kbp.

Finally, an all vs all BLASTP search was performed using the sequences included in the full visual opsin alignment (Fig. 2a; Figs. S4-S8) both as query and database. The output was filtered by subtype both among the query and subject and plotted as violin plots (Fig. S9).

All multiple sequence alignments and phylogenetic tree inference files have been deposited in FigShare (DOI: 10.6084/m9.figshare.29378324).

***Gene family classification and selection***
In order to group vertebrate genes into gene families for our analysis of conservation of synteny, we first downloaded the proteomes of vertebrates representing the major clades as well as a selection of outgroup species. The selected species and annotations from the corresponding genome assemblies were: acorn worm (*Saccoglossus kowalevskii*, Skow_1.1), amphioxus (*Branchiostoma floridae*, Bfl_VNyyK), vase tunicate (*Ciona intestinalis*, KH), pouched lamprey (*Geotria australis*), short-headed lamprey (*Mordacia mordax*), sea lamprey (*Petromyzon marinus*, kPetMar1.pri), Australian ghost shark (*Callorhincus milii*, Callorhinchus_milii-6.1.3), human (*Homo sapiens*, GRCh38), chicken (*Gallus gallus*, GRCg6a), spotted gar (*Lepisosteus oculatus*, LepOcu1), zebrafish (*Danio rerio*, GRCz11) and Japanese medaka (*Oryzias latipes*, ASM223467v1). For each proteome, only the longest peptide sequence per gene was selected using a custom script. The resulting fasta files from each species were then used as input for OrthoFinder (Emms & Kelly 2019) for gene family classification (also called orthogroups). The following setting were used for this analysis:s -M msa. The identified families were then filtered to keep only families likely to have expanded in early vertebrate whole genome duplications. This was done by identifying families with one member in acorn worm, amphioxus or vase tunicate and more than two members in either sea lamprey, pouched lamprey, short-headed lamprey, human, spotted gar, or chicken. In total we identified 1709 families with this method. Out of these families we selected those that had genes on at least two of the three chromosomes that carry visual opsin genes in the sea lamprey genome. This resulted in the identification of a total of 15 gene families (CRY, TIGD, CAMK, ATP2B, FAM3, TMCC, RAF, KDM5, PRK, LRIG, PHF, NET/ARHGEF, NT5DC, GNL, ZCCHC). Some of these families were excluded due to multiple short sequences and/or complex tree topologies (CRY, TIGD, NT5DC, ZCCHC) that indicated other modes of duplication in addition to the early vertebrate WGD events that would preclude

reliable concatenation. Out of the remaining families, those with members on at least three chromosomes in lampreys were used in further analyses (CAMK, ATP2B, TMCC, RAF, KDM5, PRK, NET/ARHGEF). For more details, see Supplementary table 4.

***Analyses of neighboring gene families***
The amino acid sequences from each gene family identified in the previous section (CAMK, ATP2B, TMCC, RAF, KDM5, PRK, NET/ARHGEF) were used as queries in TBLASTN searches against all lamprey genomes to identify members of the selected families in other lamprey genomes as well as any potential unannotated members in the pouched and short-headed lamprey genomes we performed TBLASTN searches in the genome assemblies of the sea lamprey, arctic lamprey, far eastern brook lamprey, pacific lamprey, pouched lamprey, and short-headed lamprey. The top five sequence hits for each sequence and high scoring pairs (HSPs) on these sequences located less or equal to 50 Kbp apart were merged into bed files. This distance was selected since we consider it to be a reasonable intron length to avoid merging different gene hits. FASTA sequences for each unique group of merged HSPs were extracted using bedtools. Subsequently genes were predicted on the extracted sequences using GeneWise as described for the visual opsins. For each predicted gene we extracted the longest peptide sequence and combined these with the  vase tunicate, amphioxus, and acorn worm sequences from the original family alignment generated by OrthoFinder. The resulting alignments were aligned using ClustalO with standard settings. Phylogenetic trees for inspection of families were constructed using the same settings as mentioned in the ‘Gene searches and phylogenetic analyses’ section except -bnni was not used. The resulting alignments and trees were inspected for inconsistencies. Sequences with erroneous stretches were edited either by introducing gaps or manual prediction based on sequence homology to other vertebrate sequences.

***Paralogon evolution***
The resulting alignments of the neighboring gene families as well as alignments of lamprey GNAI and PP sequences were inspected for sequences located on contigs. The identified sequences were renamed to the most likely true chromosome based on the neighboring family trees (see Supplementary table 2). All separate alignments (nine families in total) were concatenated (missing genes were coded as missing in the final alignment) using MEGA11 (Tamura et al. 2021) with standard settings. The resulting alignment was subjected to phylogenetic analysis and visualization using the same settings as mentioned in the ‘Gene searches and phylogenetic analyses’ section. For final alignment and tree inference files see FigShare (DOI: 10.6084/m9.figshare.29378324).

**Supplementary References**
Altschul SF, Gish W, Miller W, Myers EW, Lipman DJ. 1990. Basic local alignment search tool. J Mol Biol. 215:403–10. doi: 10.1016/S0022-2836(05)80360-2.
Birney E, Clamp M, Durbin R. 2004. GeneWise and Genomewise. Genome Res. 14:988–995. doi: 10.1101/gr.1865504.
Emms DM, Kelly S. 2019. OrthoFinder: phylogenetic orthology inference for comparative genomics. Genome Biol. 20:238. doi: 10.1186/s13059-019-1832-y.
Hara Y et al. 2015. Optimizing and benchmarking de novo transcriptome sequencing: From library preparation to assembly evaluation. BMC Genomics. 16. doi: 10.1186/s12864-015-2007-1.
Huang N, Li H. 2023. compleasm: a faster and more accurate reimplementation of BUSCO. Bioinformatics. 39. doi: 10.1093/bioinformatics/btad595.
Kalyaanamoorthy S, Minh BQ, Wong TKF, Von Haeseler A, Jermiin LS. 2017. ModelFinder: Fast model selection for accurate phylogenetic estimates. Nat Methods. 14:587–589. doi: 10.1038/nmeth.4285.
Lagman D et al. 2013. The vertebrate ancestral repertoire of visual opsins, transducin alpha subunits and oxytocin/vasopressin receptors was established by duplication of their shared genomic region in the two rounds of early vertebrate genome duplications. BMC Evol Biol. 13:238. doi: 10.1186/1471-2148-13-238.

Lagman D, Sundström G, Ocampo Daza D, Abalo XM, Larhammar D. 2012. Expansion of Transducin Subunit Gene Families in Early Vertebrate Tetraploidizations. Genomics. 100:203–211. doi: 10.1016/j.ygeno.2012.07.005.
Letunic I, Bork P. 2021. Interactive tree of life (iTOL) v5: An online tool for phylogenetic tree display and annotation. Nucleic Acids Res. 49:W293–W296. doi: 10.1093/nar/gkab301.
Li W, Godzik A. 2006. Cd-hit: A fast program for clustering and comparing large sets of protein or nucleotide sequences. Bioinformatics. 22:1658–1659. doi: 10.1093/bioinformatics/btl158.
Manni M, Berkeley MR, Seppey M, Zdobnov EM. 2021. BUSCO: Assessing Genomic Data Quality and Beyond. Curr Protoc. 1. doi: 10.1002/cpz1.323.
Marlétaz F et al. 2024. The hagfish genome and the evolution of vertebrates. Nature. 627:811–820. doi: 10.1038/s41586-024-07070-3.
Paradis E, Claude J, Strimmer K. 2004. APE: Analyses of phylogenetics and evolution in R language. Bioinformatics. 20:289–290. doi: 10.1093/bioinformatics/btg412.
Quinlan AR, Hall IM. 2010. BEDTools: A flexible suite of utilities for comparing genomic features. Bioinformatics. 26:841–842. doi: 10.1093/bioinformatics/btq033.
Sievers F et al. 2011. Fast, scalable generation of high-quality protein multiple sequence alignments using Clustal Omega. Mol Syst Biol. 7. doi: 10.1038/msb.2011.75.
Tamura K, Stecher G, Kumar S. 2021. MEGA11: Molecular Evolutionary Genetics Analysis Version 11. Mol Biol Evol. 38:3022–3027. doi: 10.1093/molbev/msab120.
Thi Hoang D et al. 2017. UFBoot2: Improving the Ultrafast Bootstrap Approximation. Mol. Biol. Evol. 35:518–522. doi: 10.5281/zenodo.854445.
Wang LG et al. 2020. Treeio: An R Package for Phylogenetic Tree Input and Output with Richly Annotated and Associated Data. Mol Biol Evol. 37:599–603. doi: 10.1093/molbev/msz240.
Wong T et al. 2025. IQ-TREE 3: Phylogenomic Inference Software using Complex Evolutionary Models. doi: 10.32942/X2P62N.
Xu S et al. 2022. Ggtree: A serialized data object for visualization of a phylogenetic tree and annotation data. iMeta. 1. doi: 10.1002/imt2.56.

#### Supplementary Figure Legends

**Figure S1. Phylogenetic inference of visual and non-visual opsins in jawless vertebrate genomes:** Initial phylogenetic analysis of visual opsins and closely related non-visual opsins. Clades are labeled with the name of the closest jawed vertebrate opsin subtype. Red circles indicate nodes with high-confidence support (≥80 aLRT / ≥95 UFBoot). Leaf labels include the common species name, followed by the gene name or scaffold/chromosome accession number and gene position for sequences predicted in this study. The tree is rooted with the human OPN3 sequence. See Supplementary table 3 for positions and orientation of jawless vertebrate visual opsins. The alignment with all sequences and all tree inference files are available at FigShare (DOI: 10.6084/m9.figshare.29378324).

**Figure S2. Completeness scores calculated using BUSCO and compleasm for genomes used for exhaustive visual and non-visual opsin searches:** Jawless vertebrate genomes exhibited low scores but they are regarded as underestimates caused by insufficient sensitivity in the search of exons of high sequence divergence (Hara et al. 2015). Asterisks indicate where compleasm were used when BUSCO failed in finishing the analysis.

**Figure S3. Phylogenetic inference on a reduced alignment of jawed and jawless vertebrate visual opsins:** Phylogenetic analysis of visual opsins identified in lampreys and vertebrate RefSeq proteins. The unexpected placement of the *Petromyzon marinus*, *Letheneron reissneri* and *Lethenteron camtschaticum SWS2* pseudogene sequences in this phylogeny is most likely due to their short length. Clades are labeled with the name of the closest jawed vertebrate opsin subtype at the centre of the clade. Filled red circles indicate nodes with high-confidence support (≥80 aLRT / ≥95 UFBoot). Leaf labels for RefSeq proteins include the species name, gene ID, and protein accession number. For predicted lamprey and hagfish sequences, labels include the species name, scaffold/chromosome accession number, and gene position. Leaf labels are color-coded by vertebrate group, consistent with Fig. 2a: green (teleosts), red (gars, bowfins, sturgeons, paddlefishes), orange (bichirs), brown (jawless vertebrates), blue (lobe-finned fish), and purple (cartilaginous fishes). The tree is rooted with OPN3 sequences. See Supplementary table 2 for sequence details. The alignment with all sequences and all tree inference files are available at FigShare (DOI: 10.6084/m9.figshare.29378324).

**Figures S4–S8. Subtrees of opsin gene clades (LWS, SWS2, SWS1, RH2, RH1, respectively):** Each subtree is derived from the full phylogenetic tree presented in Fig. 2a. The unexpected placement of the *Petromyzon marinus*, *Letheneron reissneri* and *Lethenteron camtschaticum* SWS2 pseudogene sequences in the full phylogeny (Fig. 2a) is most likely due to their short length. Leaf labels indicate species name, scaffold/chromosome accession number, and gene position for predicted lamprey and hagfish sequences. Leaf labels are color-coded by vertebrate group, consistent with Fig. 2a: green (teleosts), red (gars, bowfins, sturgeons, paddlefishes), orange (bichirs), brown (jawless vertebrates), blue (lobe-finned fish), and purple (cartilaginous fishes). The full phylogeny was inferred using IQ-TREE with a constraint based on a reduced taxon set. Additional sequences were added and aligned with ClustalO prior to final inference. Node support values are omitted due to limited interpretability under the constrained analysis, particularly for newly added sequences. See Supplementary table 2 for sequence details. The full alignment with all sequences and all tree inference files are available at FigShare (DOI: 10.6084/m9.figshare.29378324).

**Figure S9. All-vs-all BLASTP sequence identities within visual opsin subtypes in vertebrates:** Violin and box plots showing the distribution of pairwise percent identities among opsin protein sequences within each subtype. For each opsin subtype (LWS, SWS1, SWS2, RH2, RH1), all pairwise BLASTP percent identities between sequences of the same subtype are plotted. The violin plots illustrate the density of percent identity values, while the box plots indicate the median and interquartile range. Only comparisons between different sequences (self-hits excluded) and within the same subtype are shown.

**Figure S10. Phylogenetic inference on the concatenated alignment:** Expanded version of the phylogeny shown in Fig. 3a, maintaining the same branching order. Leaf labels include species names followed by chromosomal sequence accession numbers. Red circles indicate nodes with high-confidence support (≥80 aLRT / ≥95 UFBoot). The tree was rooted with the concatenated acorn worm (*Saccoglossus kowalevskii*) sequence. Details on the gene families included in the concatenated alignment, along with relevant comments, are provided in Supplementary table 4. The alignment with all sequences and all tree inference files are available at FigShare (DOI: 10.6084/m9.figshare.29378324).

**Figure S11. Updated phylogenetic inference of GNAT sequences in lampreys:** Expanded version of the phylogeny shown in Fig. 3b, maintaining the same branching order. Leaf labels include species common names followed by chromosomal sequence accession numbers. Red circles indicate nodes with high-confidence support (≥80 aLRT / ≥95 UFBoot). The tree was rooted with the human GNAI1-3 sequences. The alignment with all concatenated sequences and all tree inference files are available at FigShare (DOI: 10.6084/m9.figshare.29378324).

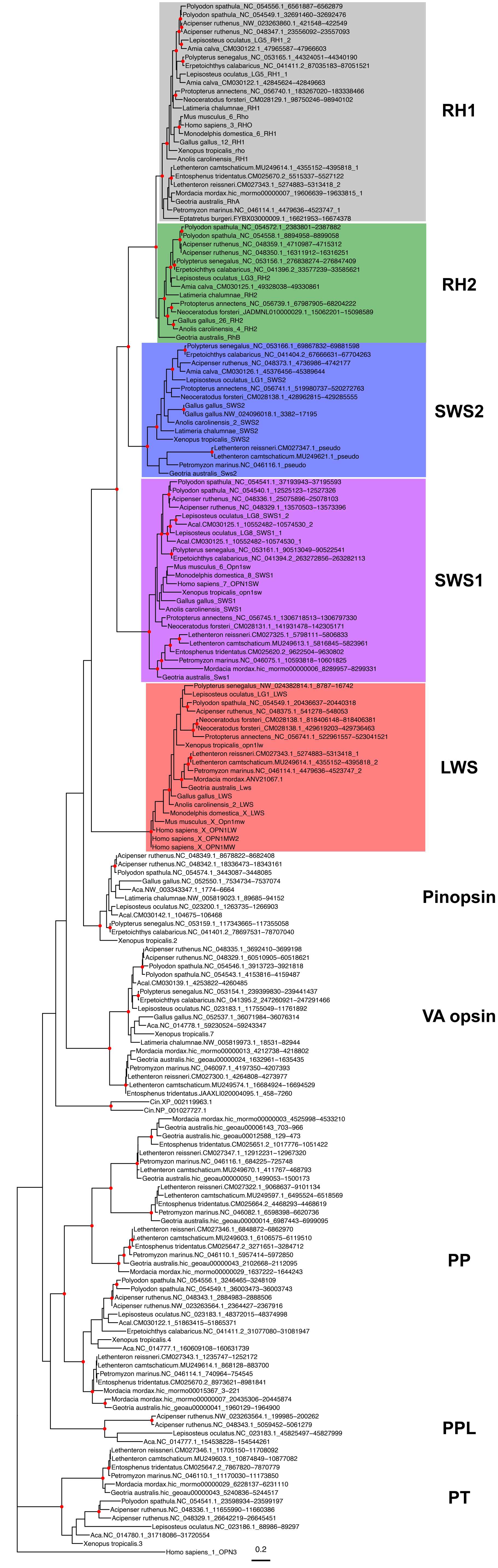
RH1
Polyodon spathula_NC_054556.1_6561887–6562879
Polyodon spathula_NC_054549.1_32691460–32692476
Acipenser ruthenus_NW_023263860.1_421548–422549
Acipenser ruthenus_NC_048347.1_23556092–23557093
Lepisosteus oculatus_LG5_RH1_2
Amia calva_CM030122.1_47965587–47966603
Polypterus senegalus_NC_053165.1_44324051–44340190
Erpetoichthys calabaricus_NC_041411.2_87035183–87051521
Lepisosteus oculatus_LG5_RH1_1
Amia calva_CM030122.1_42845624–42849663
Protopterus annectens_NC_056740.1_183267020–183338466
Neoceratodus forsteri_CM028129.1_98750246–98940102
Latimeria chalumnae_RH1
Mus musculus_6_Rho
Homo sapiens_3_RHO
Monodelphis domestica_6_RH1
Gallus gallus_12_RH1
Xenopus tropicalis_rho
Anolis carolinensis_RH1
Lethenteron camtschaticum.MU249614.1_4355152–4395818_1
Entosphenus tridentatus.CM025670.2_5515337–5527122
Lethenteron reissneri.CM027343.1_5274883–5313418_2
Mordacia mordax.hic_mormo00000007_19606639–19633815_1
Geotria australis_RhA
Petromyzon marinus.NC_046114.1_4479636–4523747_1
Eptatretus burgeri.FYBX03000009.1_16621953–16674378
RH2
Polyodon spathula_NC_054572.1_2383801–2387882
Polyodon spathula_NC_054558.1_8894958–8899058
Acipenser ruthenus_NC_048359.1_4710987–4715312
Acipenser ruthenus_NC_048350.1_16311912–16316251
Polypterus senegalus_NC_053156.1_276838274–276847409
Erpetoichthys calabaricus_NC_041396.2_33577239–33585621
Lepisosteus oculatus_LG3_RH2
Amia calva_CM030125.1_49328038–49330861
Latimeria chalumnae_RH2
Protopterus annectens_NC_056739.1_67987905–68204222
Neoceratodus forsteri_JADMNL010000029.1_15062201–15098589
Gallus gallus_26_RH2
Anolis carolinensis_4_RH2
Geotria australis_RhB
SWS2
Polypterus senegalus_NC_053166.1_69867832–69881598
Erpetoichthys calabaricus_NC_041404.2_67666631–67704263
Acipenser ruthenus_NC_048373.1_4736986–4742177
Amia calva_CM030126.1_45376456–45389644
Lepisosteus oculatus_LG1_SWS2
Protopterus annectens_NC_056741.1_519980737–520272763
Neoceratodus forsteri_CM028138.1_428962815–429285555
Gallus gallus_SWS2
Gallus gallus.NW_024096018.1_3382–17195
Anolis carolinensis_2_SWS2
Latimeria chalumnae_SWS2
Xenopus tropicalis_SWS2
Lethenteron reissneri.CM027347.1_pseudo
Lethenteron camtschaticum.MU249621.1_pseudo
Petromyzon marinus.NC_046116.1_pseudo
Geotria australis_Sws2
SWS1
Polyodon spathula_NC_054541.1_37193943–37195593
Polyodon spathula_NC_054540.1_12525123–12527326
Acipenser ruthenus_NC_048336.1_25075896–25078103
Acipenser ruthenus_NC_048329.1_13570503–13573396
Lepisosteus oculatus_LG8_SWS1_2
Acal.CM030125.1_10552482–10574530_2
Lepisosteus oculatus_LG8_SWS1_1
Acal.CM030125.1_10552482–10574530_1
Polypterus senegalus_NC_053161.1_90513049–90522541
Erpetoichthys calabaricus_NC_041394.2_263272856–263282113
Mus musculus_6_Opn1sw
Monodelphis domestica_8_SWS1
Homo sapiens_7_OPN1SW
Xenopus tropicalis_opn1sw
Gallus gallus_SWS1
Anolis carolinensis_SWS1
Protopterus annectens_NC_056745.1_1306718513–1306797330
Neoceratodus forsteri_CM028131.1_141931478–142305171
Lethenteron reissneri.CM027325.1_5798111–5806833
Lethenteron camtschaticum.MU249613.1_5816845–5823961
Entosphenus tridentatus.CM025620.2_9622504–9630802
Petromyzon marinus.NC_046075.1_10593818–10601825
Mordacia mordax.hic_mormo00000006_8289957–8299331
Geotria australis_Sws1
LWS
Polypterus senegalus_NW_024382814.1_8787–16742
Lepisosteus oculatus_LG1_LWS
Polyodon spathula_NC_054549.1_20436637–20440318
Acipenser ruthenus_NC_048375.1_541278–548053
Neoceratodus forsteri_CM028138.1_818406148–818406381
Neoceratodus forsteri_CM028138.1_429619203–429736463
Protopterus annectens_NC_056741.1_522961557–523041521
Xenopus tropicalis_opn1lw
Lethenteron reissneri.CM027343.1_5274883–5313418_1
Lethenteron camtschaticum.MU249614.1_4355152–4395818_2
Petromyzon marinus.NC_046114.1_4479636–4523747_2
Mordacia mordax.ANV21067.1
Geotria australis_Lws
Gallus gallus_LWS
Anolis carolinensis_2_LWS
Monodelphis domestica_X_LWS
Mus musculus_X_Opn1mw
Homo sapiens_X_OPN1LW
Homo sapiens_X_OPN1MW2
Homo sapiens_X_OPN1MW
Pinopsin
Acipenser ruthenus.NC_048349.1_8678822–8682408
Acipenser ruthenus.NC_048342.1_18336473–18343161
Polyodon spathula.NC_054574.1_3443087–3448085
Gallus gallus.NC_052550.1_7534734–7537074
Aca.NW_003343347.1_1774–6664
Latimeria chalumnae.NW_005819023.1_89685–94152
Lepisosteus oculatus.NC_023200.1_1263735–1266903
Acal.CM030142.1_104675–106468
Polypterus senegalus.NC_053159.1_117343665–117355058
Erpetoichthys calabaricus.NC_041401.2_78697531–78707040
Xenopus tropicalis.2
VA opsin
Acipenser ruthenus.NC_048335.1_3692410–3699198
Acipenser ruthenus.NC_048329.1_60510905–60518621
Polyodon spathula.NC_054546.1_3913723–3921818
Polyodon spathula.NC_054543.1_4153816–4159487
Acal.CM030139.1_4253822–4260485
Polypterus senegalus.NC_053154.1_239399830–239441437
Erpetoichthys calabaricus.NC_041395.2_247260921–247291466
Lepisosteus oculatus.NC_023183.1_11755049–11761892
Gallus gallus.NC_052537.1_36071984–36076314
Aca.NC_014778.1_59230524–59243347
Xenopus tropicalis.7
Latimeria chalumnae.NW_005819973.1_18531–82944
Mordacia mordax.hic_mormo00000013_4212738–4218802
Geotria australis.hic_geoau00000024_1632961–1635435
Petromyzon marinus.NC_046097.1_4197350–4207393
Lethenteron reissneri.CM027300.1_4264808–4273977
Lethenteron camtschaticum.MU249574.1_16684924–16694529
Entosphenus tridentatus.JAAXLI020004095.1_458–7260
Cin.XP_002119963.1
Cin.NP_001027727.1
PP
Mordacia mordax.hic_mormo00000003_4525998–4533210
Geotria australis.hic_geoau00006143_703–966
Geotria australis.hic_geoau00012588_129–473
Entosphenus tridentatus.CM025651.2_1017776–1051422
Lethenteron reissneri.CM027347.1_12912231–12967320
Petromyzon marinus.NC_046116.1_684225–725748
Lethenteron camtschaticum.MU249670.1_411767–468793
Geotria australis.hic_geoau00000050_1499053–1500173
Lethenteron reissneri.CM027322.1_9068637–9101134
Lethenteron camtschaticum.MU249597.1_6495524–6518569
Entosphenus tridentatus.CM025664.2_4468293–4468619
Petromyzon marinus.NC_046082.1_6598398–6620736
Geotria australis.hic_geoau00000014_6987443–6999095
Lethenteron reissneri.CM027346.1_6848872–6862970
Lethenteron camtschaticum.MU249603.1_6106575–6119510
Entosphenus tridentatus.CM025647.2_3271651–3284712
Petromyzon marinus.NC_046110.1_5957414–5972850
Geotria australis.hic_geoau00000043_2102668–2112095
Mordacia mordax.hic_mormo00000029_1637222–1644243
Polyodon spathula.NC_054556.1_3246465–3248109
Polyodon spathula.NC_054549.1_36003473–36003747
Acipenser ruthenus.NC_048343.1_2884983–2888506
Acipenser ruthenus.NW_023263564.1_2364427–2367916
Lepisosteus oculatus.NC_023183.1_48372015–48374998
Acal.CM030122.1_51863415–51865371
Erpetoichthys calabaricus.NC_041411.2_31077080–31081947
Xenopus tropicalis.4
Aca.NC_014777.1_160609108–160631739
Lethenteron reissneri.CM027343.1_1235747–1252172
Lethenteron camtschaticum.MU249614.1_868128–883700
Petromyzon marinus.NC_046114.1_740964–754545
Entosphenus tridentatus.CM025670.2_8973621–8981841
Mordacia mordax.hic_mormo00015367_3–221
Mordacia mordax.hic_mormo00000007_20435306–20445874
Geotria australis.hic_geoau00000041_1960129–1964900
PPL
Acipenser ruthenus.NW_023263564.1_199985–200262
Acipenser ruthenus.NC_048343.1_5059452–5061279
Lepisosteus oculatus.NC_023183.1_45825497–45827999
Aca.NC_014777.1_154538228–154544261
PT
Lethenteron reissneri.CM027346.1_11705150–11708092
Lethenteron camtschaticum.MU249603.1_10874849–10877082
Entosphenus tridentatus.CM025647.2_7867820–7870779
Petromyzon marinus.NC_046110.1_11170030–11173850
Mordacia mordax.hic_mormo00000029_6228137–6231110
Geotria australis.hic_geoau00000043_5240836–5244517
Polyodon spathula.NC_054541.1_23598934–23599197
Acipenser ruthenus.NC_048336.1_11655990–11660386
Acipenser ruthenus.NC_048329.1_26642219–26645451
Lepisosteus oculatus.NC_023186.1_89886–89297
Aca.NC_014780.1_31718086–31720554
Xenopus tropicalis.3
Homo sapiens_1_OPN3
0.2

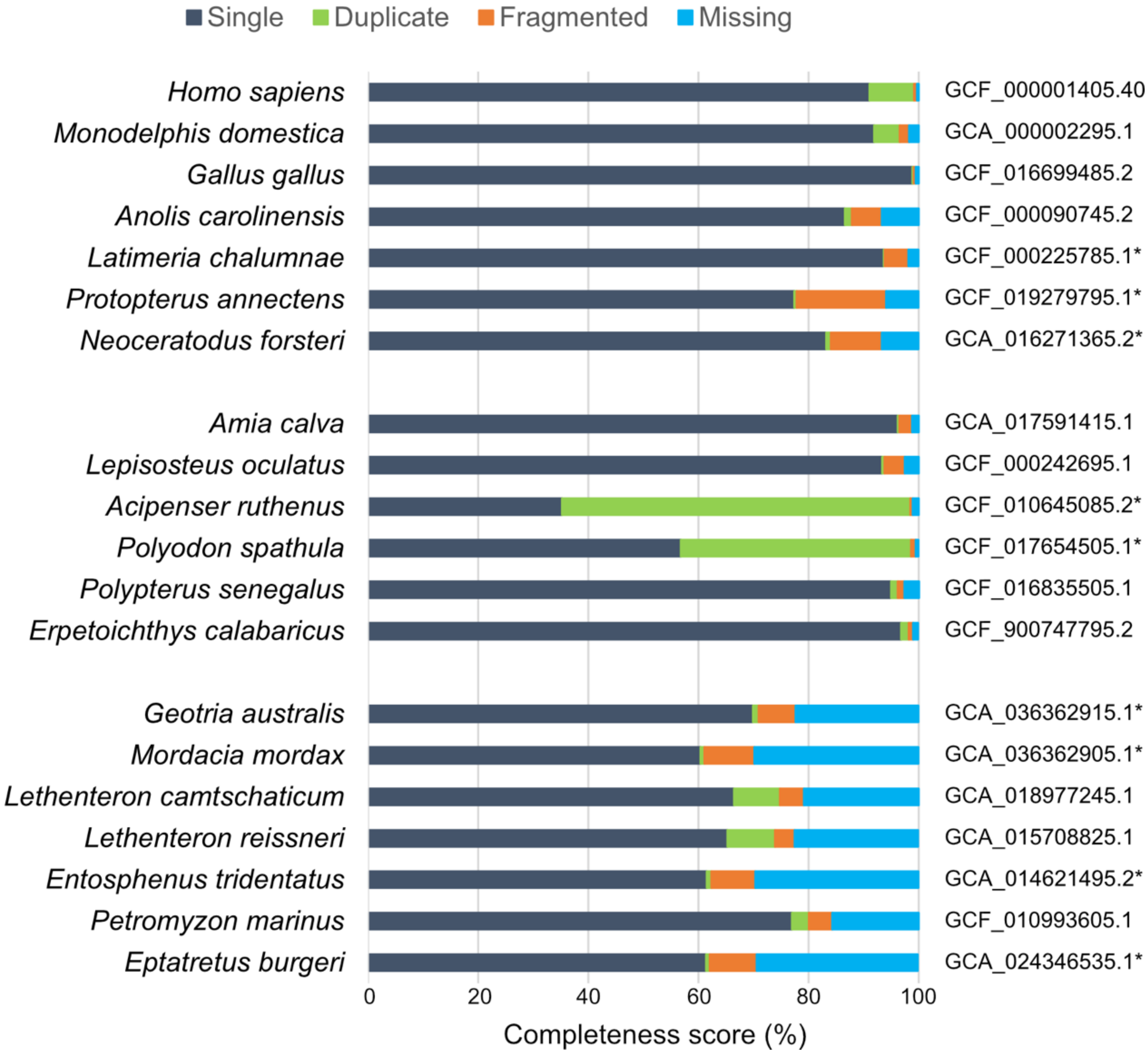

Single
Duplicate
Fragmented
Missing
Homo sapiens
Monodelphis domestica
Gallus gallus
Anolis carolinensis
Latimeria chalumnae
Protopterus annectens
Neoceratodus forsteri
Amia calva
Lepisosteus oculatus
Acipenser ruthenus
Polyodon spathula
Polypterus senegalus
Erpetoichthys calabaricus
Geotria australis
Mordacia mordax
Lethenteron camtschaticum
Lethenteron reissneri
Entosphenus tridentatus
Petromyzon marinus
Eptatretus burgeri
GCF_000001405.40
GCA_000002295.1
GCF_016699485.2
GCF_000090745.2
GCF_000225785.1*
GCF_019279795.1*
GCA_016271365.2*
GCA_017591415.1
GCF_000242695.1
GCF_010645085.2*
GCF_017654505.1*
GCF_016835505.1
GCF_900747795.2
GCA_036362915.1*
GCA_036362905.1*
GCA_018977245.1
GCA_015708825.1
GCA_014621495.2*
GCF_010993605.1
GCA_024346535.1*
0
20
40
60
80
100
Completeness score (%)

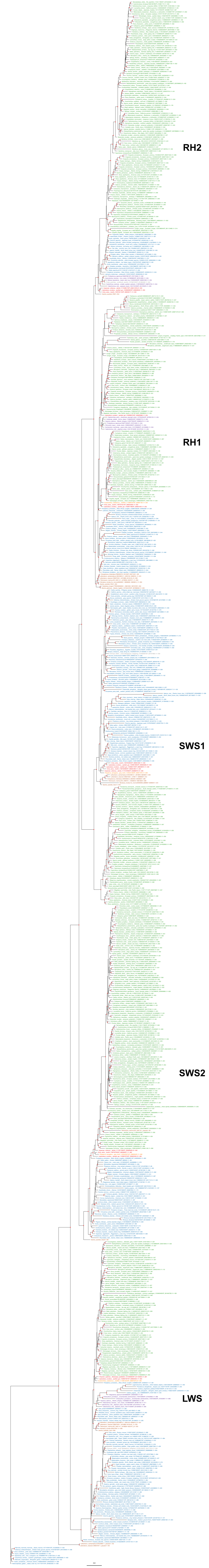
RH2
RH1
SWS1
SWS2
LWS

Opsin Clade: LWS

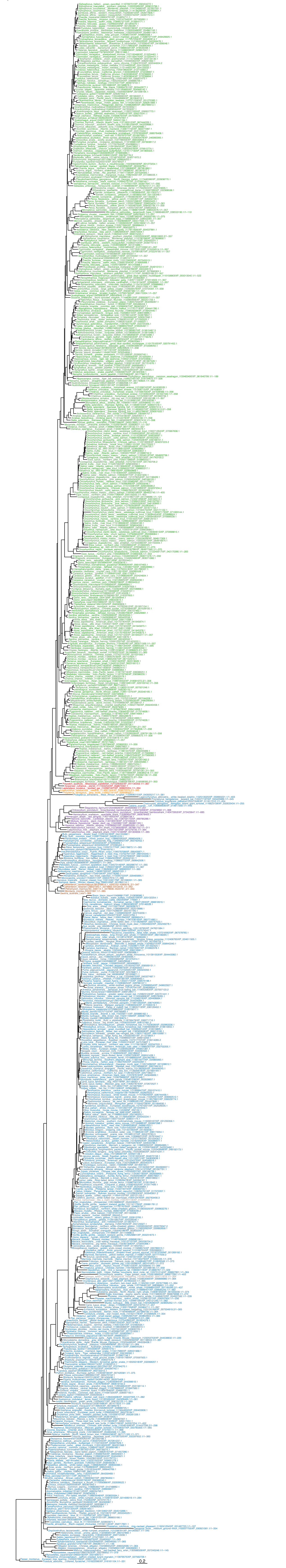

Opsin Clade: SWS2

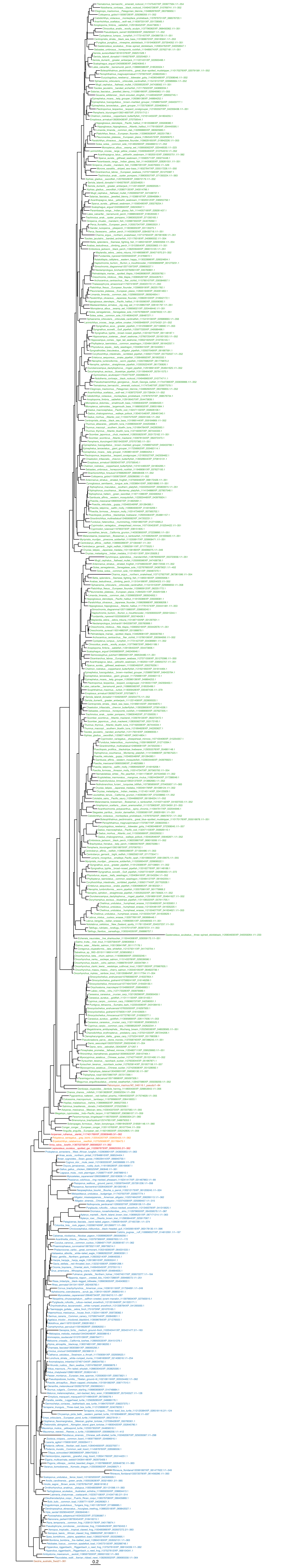

#### Opsin Clade: SWS1

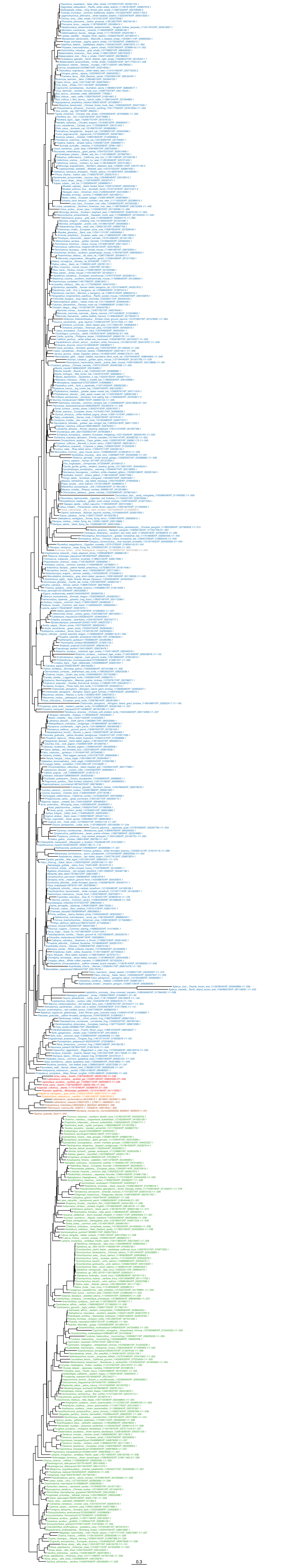

Opsin Clade: RH2

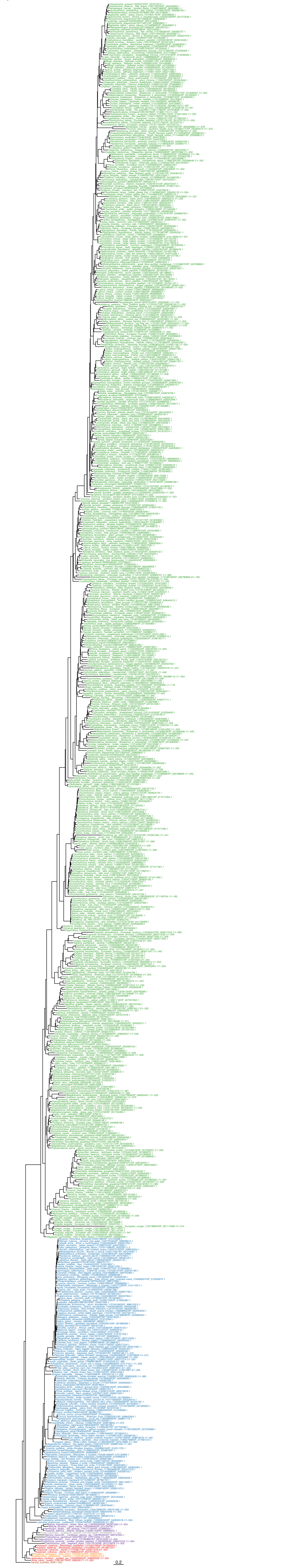

Opsin Clade: Rho

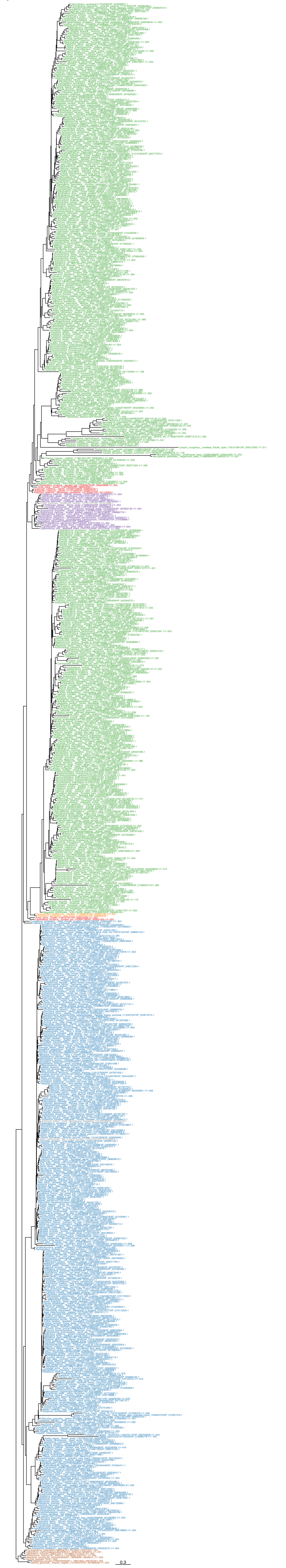

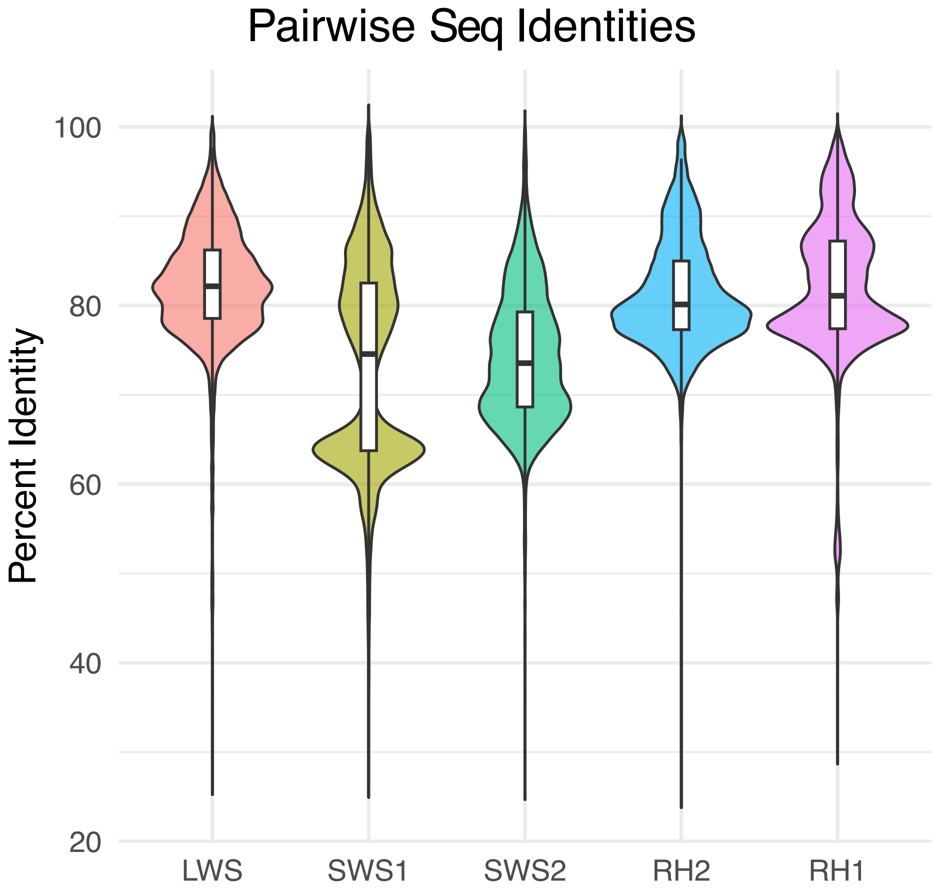
Pairwise Seq Identities
Percent Identity
100
80
60
40
20
LWS
SWS1
SWS2
RH2
RH1

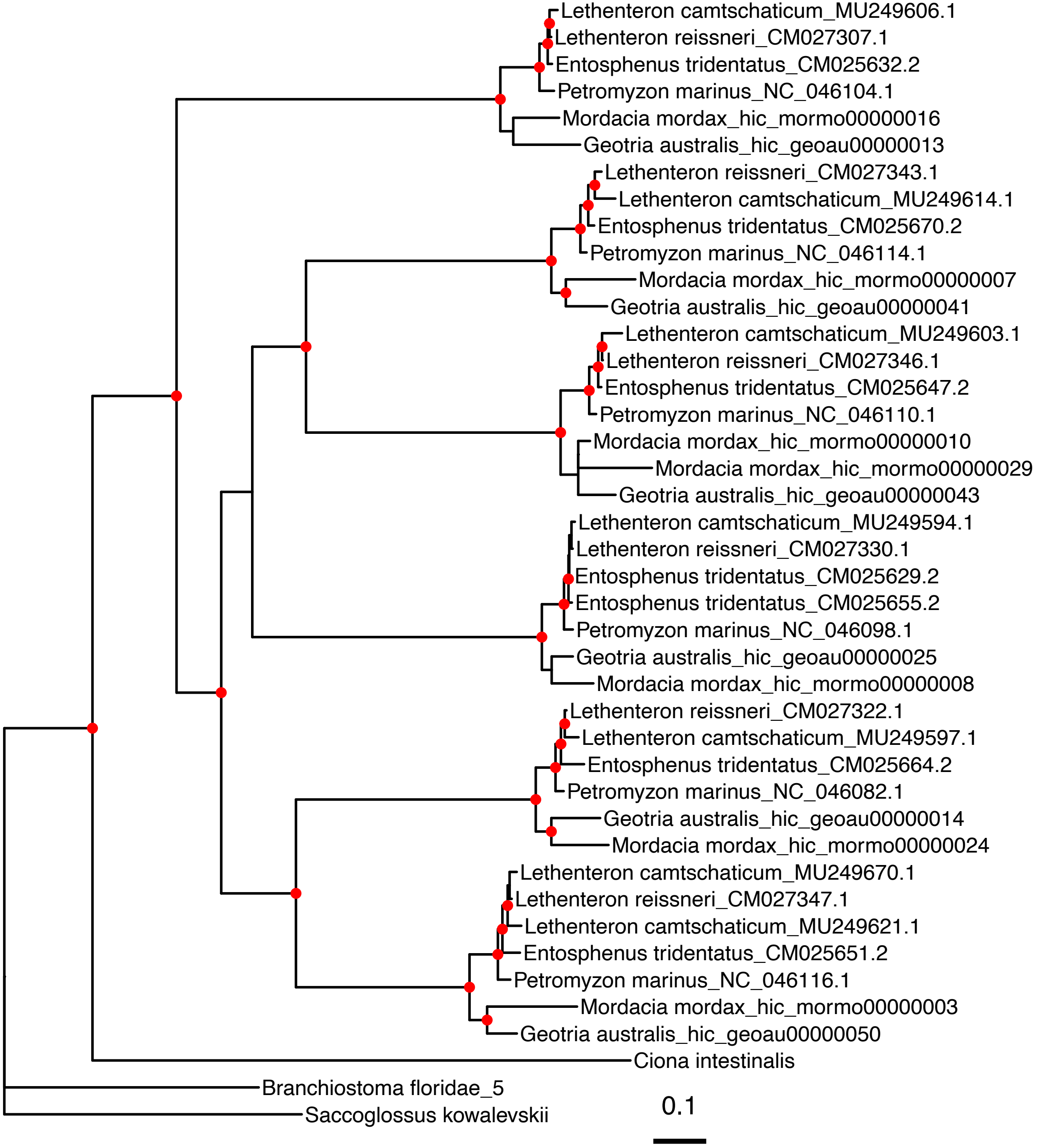

Lethenteron camtschaticum_MU249606.1
Lethenteron reissneri_CM027307.1
Entosphenus tridentatus_CM025632.2
Petromyzon marinus_NC_046104.1
Mordacia mordax_hic_mormo00000016
Geotria australis_hic_geoau00000013
Lethenteron reissneri_CM027343.1
Lethenteron camtschaticum_MU249614.1
Entosphenus tridentatus_CM025670.2
Petromyzon marinus_NC_046114.1
Mordacia mordax_hic_mormo00000007
Geotria australis_hic_geoau00000041
Lethenteron camtschaticum_MU249603.1
Lethenteron reissneri_CM027346.1
Entosphenus tridentatus_CM025647.2
Petromyzon marinus_NC_046110.1
Mordacia mordax_hic_mormo00000010
Mordacia mordax_hic_mormo00000029
Geotria australis_hic_geoau00000043
Lethenteron camtschaticum_MU249594.1
Lethenteron reissneri_CM027330.1
Entosphenus tridentatus_CM025629.2
Entosphenus tridentatus_CM025655.2
Petromyzon marinus_NC_046098.1
Geotria australis_hic_geoau00000025
Mordacia mordax_hic_mormo00000008
Lethenteron reissneri_CM027322.1
Lethenteron camtschaticum_MU249597.1
Entosphenus tridentatus_CM025664.2
Petromyzon marinus_NC_046082.1
Geotria australis_hic_geoau00000014
Mordacia mordax_hic_mormo00000024
Lethenteron camtschaticum_MU249670.1
Lethenteron reissneri_CM027347.1
Lethenteron camtschaticum_MU249621.1
Entosphenus tridentatus_CM025651.2
Petromyzon marinus_NC_046116.1
Mordacia mordax_hic_mormo00000003
Geotria australis_hic_geoau00000050
Ciona intestinalis
Branchiostoma floridae_5
Saccoglossus kowalevskii
0.1

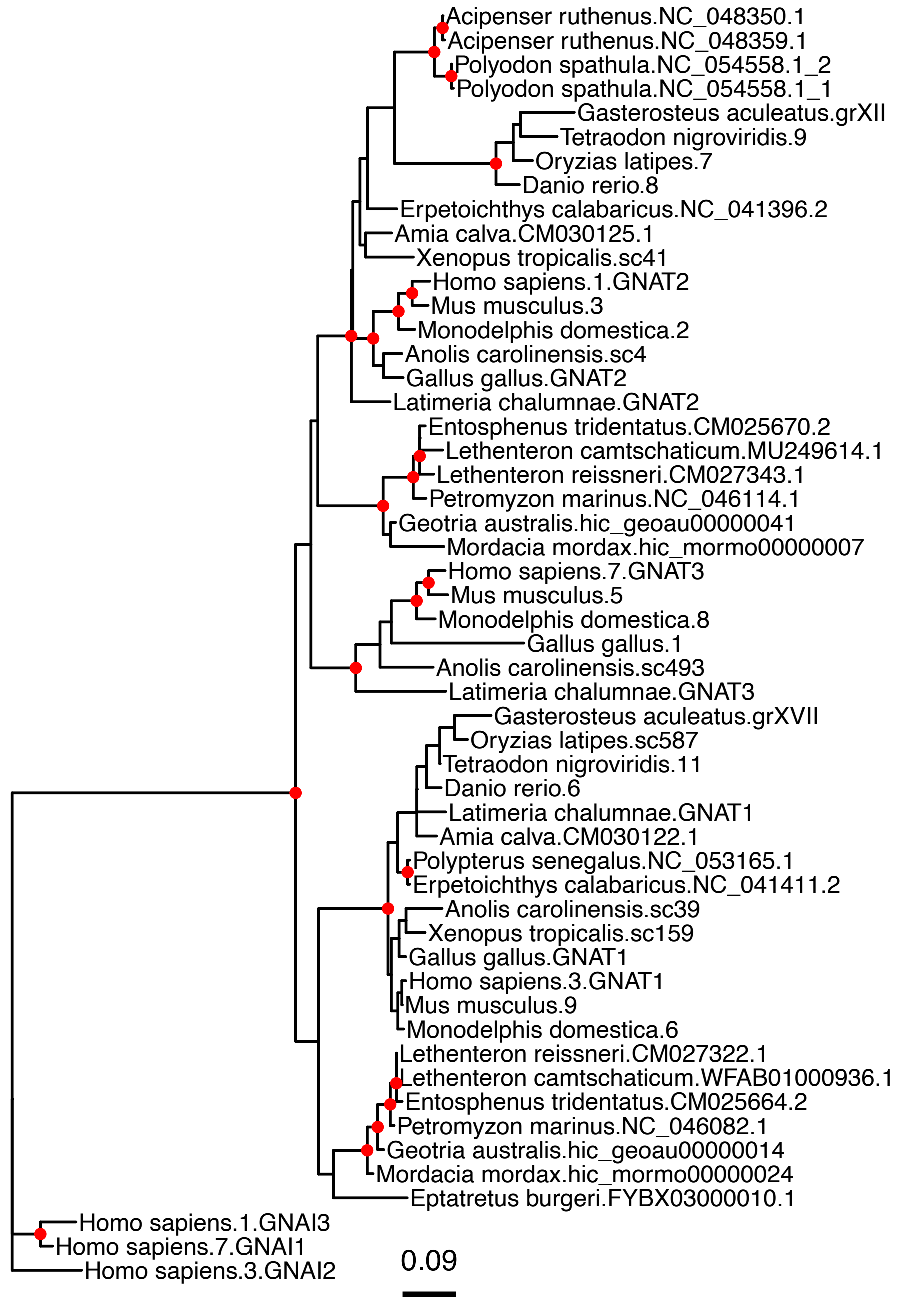
Acipenser ruthenus.NC_048350.1
Acipenser ruthenus.NC_048359.1
Polyodon spathula.NC_054558.1_2
Polyodon spathula.NC_054558.1_1
Gasterosteus aculeatus.grXII
Tetraodon nigroviridis.9
Oryzias latipes.7
Danio rerio.8
Erpetoichthys calabaricus.NC_041396.2
Amia calva.CM030125.1
Xenopus tropicalis.sc41
Homo sapiens.1.GNAT2
Mus musculus.3
Monodelphis domestica.2
Anolis carolinensis.sc4
Gallus gallus.GNAT2
Latimeria chalumnae.GNAT2
Entosphenus tridentatus.CM025670.2
Lethenteron camtschaticum.MU249614.1
Lethenteron reissneri.CM027343.1
Petromyzon marinus.NC_046114.1
Geotria australis.hic_geoau00000041
Mordacia mordax.hic_mormo00000007
Homo sapiens.7.GNAT3
Mus musculus.5
Monodelphis domestica.8
Gallus gallus.1
Anolis carolinensis.sc493
Latimeria chalumnae.GNAT3
Gasterosteus aculeatus.grXVII
Oryzias latipes.sc587
Tetraodon nigroviridis.11
Danio rerio.6
Latimeria chalumnae.GNAT1
Amia calva.CM030122.1
Polypterus senegalus.NC_053165.1
Erpetoichthys calabaricus.NC_041411.2
Anolis carolinensis.sc39
Xenopus tropicalis.sc159
Gallus gallus.GNAT1
Homo sapiens.3.GNAT1
Mus musculus.9
Monodelphis domestica.6
Lethenteron reissneri.CM027322.1
Lethenteron camtschaticum.WFAB01000936.1
Entosphenus tridentatus.CM025664.2
Petromyzon marinus.NC_046082.1
Geotria australis.hic_geoau00000014
Mordacia mordax.hic_mormo00000024
Eptatretus burgeri.FYBX03000010.1
Homo sapiens.1.GNAI3
Homo sapiens.7.GNAI1
Homo sapiens.3.GNAI2
0.09
